# Dynamin-2 controls complement receptor 3- mediated phagocytosis completion and closure

**DOI:** 10.1101/2021.02.14.431164

**Authors:** Anna Mularski, Ryszard Wimmer, Floriane Arbaretaz, Gabriel Le Goff, Florence Niedergang

## Abstract

Phagocytosis is the mechanism of the internalization of large particles, microorganisms and cellular debris. The complement pathway represents one of the first mechanisms of defense against infection and the complement receptor 3 (CR3), which is highly expressed on macrophages, is a major receptor for many pathogens and debris. Key to dissecting the mechanisms by which CR3-mediated phagocytosis occurs, is understanding how the complex actin binding protein machinery and associated regulators interact with actin during phagocytosis, from triggering of receptor, through to phagosome formation and closure. However, how CR3-mediated phagosome completion and closure are orchestrated is not known. Here, we reveal that dynamin-2 is recruited concomitantly with polymerised actin at the site of the nascent phagosomes and accumulates until membrane scission. Inhibition of dynamin activity leads to stalled phagocytic cups and a decrease in the amount of F-actin at the site of phagocytosis. Acute inhibition of dynamin activity in living phagocytosing cells established that dynamin-2 plays a critical role in the effective scission of the CR3-phagosome from the plasma membrane. Thus, dynamin-2 has two distinct roles in CR3-mediated phagocytosis, in the assembly of the F-actin phagocytic cup and during phagosome scission.

## Introduction

Phagocytosis is the mechanism of the internalization of large particles, microorganisms and cellular debris (Niedergang and Grinstein, 2018; Uribe-Querol and Rosales, 2020). In the professional phagocytes of the immune system, phagocytosis plays a key role in the development of immune responses via cytokine secretion and presentation of antigen derived from the internalized material. Phagocytosis is also important for normal turnover and remodelling of tissues and disposal of dead cells. Phagocytosis is initiated by the triggering of surface receptors that bind directly to surface determinants of microorganisms or receptors for opsonins like immunoglobulins or complement that coat the particulate antigen. Signal transduction downstream of the phagocytic receptors involves local reorganization of the plasma membrane, and the underlying actin cytoskeleton, resulting in the internalisation of the phagocytic target.

The complement receptor (CR) 3, which is highly expressed on macrophages, is a direct receptor for many pathogens and binds activated complement molecules deposited on microorganisms or cells (Torres-Gomez et al., 2020a). It is a phagocytic integrin also known as αMβ2 that needs to be activated to bind to extracellular ligands. The actin dynamics play a key role both in the activation of the integrin receptors and in the downstream signaling pathways that are controlled by the small GTPase RhoA, the formin mDia1 and the Arp2/3 nucleator (Caron and Hall, 1998; Colucci-Guyon et al., 2005; May et al., 2000). In addition, a subtle interplay between the microtubules and the actin cytoskeleton is important for CR3-mediated phagocytosis (Allen and Aderem, 1996; Binker et al., 2007; Lewkowicz et al., 2008). The early steps of CR3 engagement and signalling have been dissected to show how the Syk kinase and focal adhesion proteins act as a molecular clutch to promote phagocytosis (Jaumouille et al., 2019). However, much less is known about the later steps of phagosome formation and closure downstream of the CR3. CR3-mediated phagocytosis was initially reported to rely on particle “sinking”, without extensive formation of membranous lamellipodia (Allen and Aderem, 1996; Kaplan et al., 1977). Some membrane extension and ruffling activity has however been associated with CR3–mediated phagocytosis (Jaumouille et al., 2019; Patel and Harrison, 2008).

Here we investigated whether dynamin-2, an important mediator of budding and scission of small vesicles, that also has the property to bind and bundle actin filaments, plays a role in integrin-phagosome formation and scission. Complement-mediated phagocytosis was investigated in primary human monocyte-derived macrophages (hMDMs) using super resolution microscopy and phagocytosis assays with actin quantification. Concomitant recruitment of F-actin and Dynamin-2 was monitored at the onset of CR3-mediated phagocytosis by live cell imaging with total internal reflection fluorescence microscopy (TIRFM). Actin quantification in phagocytosis assays revealed that Dynamin-2 is vital for the formation of the phagocytic cup, and therefore, efficient phagocytosis. Dynamic live cell imaging revealed that dynamin-2 is required for the CR3-phagosome closure in macrophages.

## Results

### Dynamin-2 is recruited concomitantly with actin at the site of CR3-phagosome formation

We used primary human macrophages and direct stochastic optical reconstruction microscopy (dSTORM) to inform our studies of the role on dynamin-2 in CR3-mediated phagocytosis. First, to study the earliest stages of CR3-phagosome formation, frustrated phagocytosis experiments (shown schematically in Figure 1A) were performed. For this, human monocyte-derived macrophages (hMDMs) were allowed to spread on coverslips opsonised first with IgM, and then with C5-deficient complement. Cells were fixed and F-actin stained, and prior to dSTORM observation of the frustrated phagocytic cup, it was verified that the cell body was visible at a higher z. On the complement opsonised coverlips, frustrated phagocytic regions were irregular in shape, and revealed no areas of reduced F-actin density (Figure 1B, lower), which is different from the reported subcellular circular distribution of F-actin upon FcR ligation (Barger et al., 2019; Marie-Anais et al., 2016b; Marion et al., 2012). To quantify the distribution of F-actin in the spreading zone, the smallest possible circle (red Figure 1B) was drawn around frustrated phagocytic cups, and 8 radial lengths (dashed yellow, Figure 1B) were measured as demonstrated in Figure 1B.

**Figure 1:**
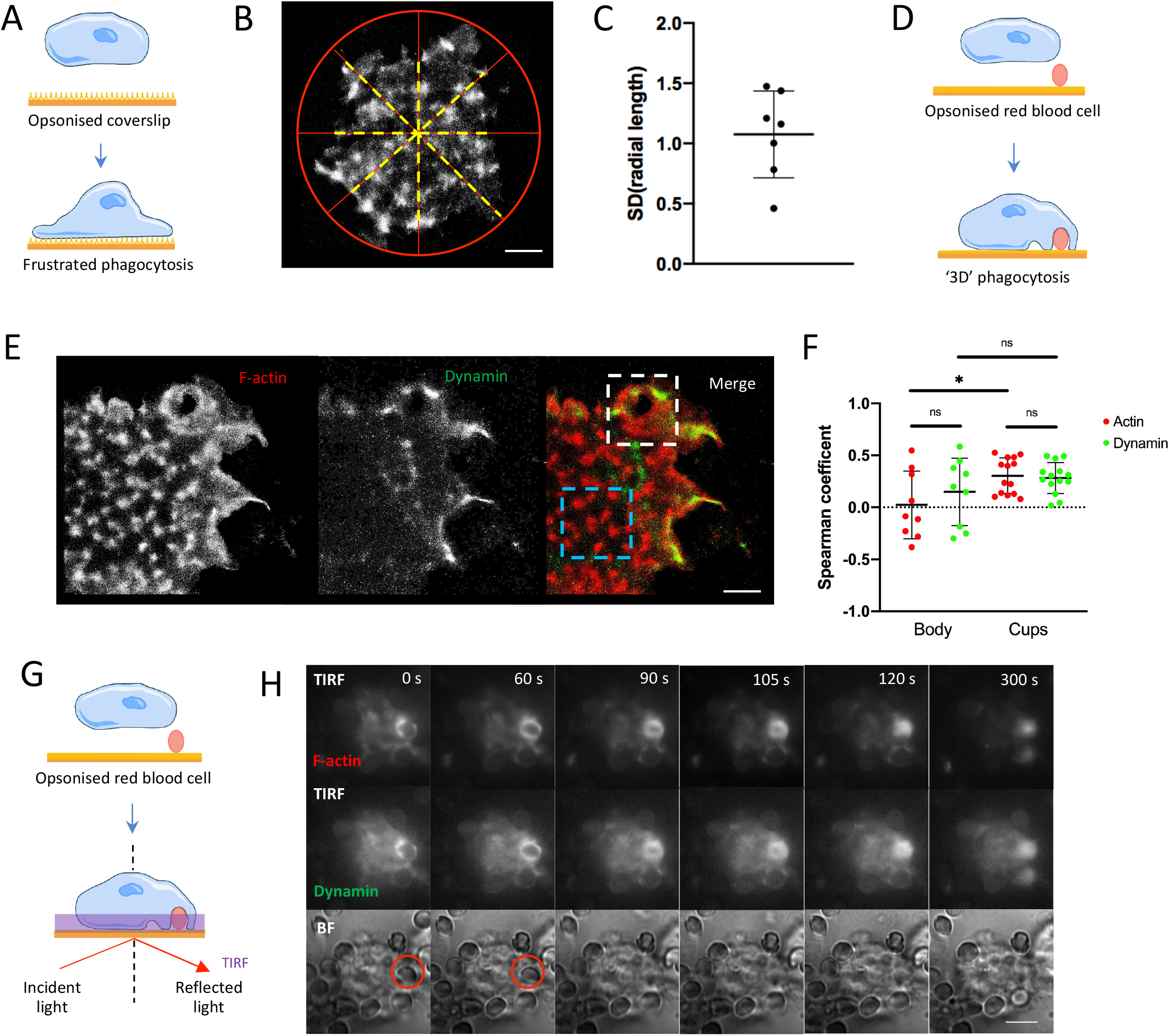
STORM visualisation of hMDMs undergoing frustrated and ‘3D’ phagocytosis show accumulation of F-actin and dynamin-2. Schematic representation of a cell undergoing frustrated phagocytosis (A). Primary human macrophages are deposited on coverslips coated with IgG (FcR) or IgM + complement (CR3) and then fixed, permeabilized and actin was labeled with phalloidin-Alexa 647 and cells were visualised with STORM (B), bar = 2.5 μm. Radial lengths of the frustrated phagocytic cups were measurered and the standard devations of these were plotted (C, 7 cells). Schematic representation of a cell undergoing ‘3D’ phagocytosis (D). Primary human macrophages are allowed to sediment on to coverslips coated with loosely bound IgG (FcR) or IgM + complement (CR3) opsonised red blood cells and then fixed, permeabilized and labeled. Actin was labelled with phalloidin-Alexa 647 and Dynamin-2 with Alexa-555. Cells were visualised with STORM (E), bar = 2.5 μm. Image analysis determined Spearman coeffcients for actin (red circles) and dynamin (green circles) in regions of interest in cell body (E, blue dashed line, 9 cell body regions) and phagocytic cups (E, white dashed line, 14 phagocytic cups) (F). Phagosome closure assay was performed using RAW 264.7 macrophages transiently expressing both Lifeact-mcherry and Dynamin-2-GFP. Shown schematically in G, cells are allowed to phagocytose opsonized particles that are non-covalently bound to the coverslip, allowing the site of closure to be observed using TIRFM. TIRFM images of Lifeact-mcherry (H, top row) and Dynamin-2-GFP (H, middle row) and brightfield images (H, bottom row) were captured every 2 s for 10 min at 37°. The RBC phagocytosed is clearly visible in brightfield images at 0 s and 60 s (B, bottom row, red circles), but pulled from the focal plane by the cell from 90s. Bar = 5 μm.

The standard deviations of these radial lengths for 7 cells undergoing frustrated phagocytosis condition were plotted (Figure 1C). By this measure, as phagocytic spreading approaches circular, the standard devation of the radial lengths would approach 0. The plot in Figure 1C indicates that the distribution of F-actin in the frustrated CR3-mediated phagocytic spreading zone was heterogeneous and did not exhibit the circularity that is characteristic of FcR-phagocytosis (Barger et al., 2019; Marie-Anais et al., 2016b; Marion et al., 2012). The closer the phagocytic spreading is to a circle, the closer the standard devation of the radial lenths would be to 0.

To view the phagosome morphology in a ‘3D’ arrangement, and at a later stage, hMDMs were allowed to perform a “3D phagosome closure assay” as described previously (Marie-Anais et al., 2016a; Mularski et al., 2018). For this, hMDMs sediment onto coverslips coated with red blood cells (RBCs) opsonised with IgM followed by complement (C’-IgM-RBC), and noncovalently bound to a poly-L-lysine activated surface (shown schematically in Figure 1D). Cells were allowed to phagocytose the opsonised RBCs for 10 – 15 minutes prior to fixation and staining to detect both F-actin and dynamin-2 by dSTORM (Figure 1E). F-actin is accumulated in focal adhesion structures as well as at the site of nascent phagosomes (Figure 1E, left panel). Dynamin-2 is enriched in these F-actin rich phagocytic cups, although not exactly at the same place as F-actin accumulations (Figure 1E, middle panel). A tessellation-based colocalization analysis (Levet et al., 2015) was performed yielding a colocalization value for each single molecule localization of each species separately, minimizing false positive colocalization. The colocalization parameter, the Spearman coefficient, can have values from -1 (anticorrelation) through 0 (no correlation and therefore a low probability of colocalisation) to 1 (perfect correlation and therefore a high probability of colocalisation). The analysis allowed for a comparison of the actin-dynamin and dynamin-actin colocalization in the cell body (blue dashed line, Figure 1E) and the phagocytic cups (white dashed line, Figure 1E). The Spearman coefficients of actin with dynamin and dynamin with actin from analysis of regions of interest in the cell body are close to zero (means of 0.024 and 0.15 respectively), indicating a low probability of colocalisation (Figure 1F). The Spearman coefficients for actin-dynamin and dynamin-actin colocalisation from regions of interest around phagocytic cups yield positive means (means of 0.30 and 0.28 respectively), indicating a higher probability of colocalization in the phagocytic cups than in the cell body (Figure 1F).

To better describe the role dynamin-2 plays in CR3-dependent phagocytosis, we used TIRFM to observe the dynamics of recruitment of F-actin and dynamin-2 in RAW264.7 macrophages transfected to transiently express actin-mCherry and dynamin-2-GFP. Cells were exposed to coverslips coated with C’-IgM-RBC) noncovalently bound to poly-L-lysine activated surface. After transfected cells were allowed to sediment to the surface, phagocytic cup extension and closure around the bound RBCs were observed in the TIRF region, as described in detail in (Marie-Anais et al., 2016a; Mularski et al., 2018) and shown schematically in Figure 1G. A time-course of a macrophage undergoing CR3 mediated phagocytosis is shown in Figure 1H. F-actin and dynamin-2 are recruited concomitantly throughout the phagocytic event, from membrane extension around the RBC, through to fusion and scission, before clearance back towards cortical levels. Dynamin-2 was recruited with actin at the tips of membrane extensions surrounding the RBC and accumulated at the site of phagosome closure and scission before clearance. These results indicate that dynamin-2 is part of the phagosome formation and completion steps.

### Phagocytosis, but not association, is impaired by dynamin-2 inhibition

To determine the role of dynamin-2 in CR3-mediated phagocytosis, we inhibited its function prior to performing phagocytosis assays. hMDMs, treated with DMSO (control) or dynasore (80 μM) were centrifuged in medium containing C’-IgM-RBCs for 2 min to synchronise phagocytosis. Macrophages were either fixed at this point to assess the efficiency of RBC association (0 min), or, were allowed to phagocytose C’IgM-RBCs for 60 min at 37°C to evaluate phagocytic efficiency. For both time points, following fixation, macrophages were stained for external RBCs (Figure 2A, B, D and E). The number of external and internal RBCs was quantified. The association index, the average number of cell-associated (bound + internalised) RBCs at 0 min and the phagocytic index, the average number of internalized cells at 60 min, were calculated. The indices of association from 3 independent experiments on different donors (Figure 2C), revealed no significant difference between cells treated with DMSO or dynasore, in their ability to engage with C’IgM-RBCs. The indices of phagocytosis (Figure 2F) however, demonstrate that phagocytic efficiency is significantly reduced in cells treated with dynasore relative to control cells (∗∗ p < 0.01). This result indicates that CR3 receptor activation at the cell surface is not affected after treatment with the dynamin-2 inhibitor. As such, inhibiting dynamin-2 with dynasore monohydrate leads to the impairment of phagocytosis without interfering with integrin-receptor binding to the complement ligand.

**Figure 2:**
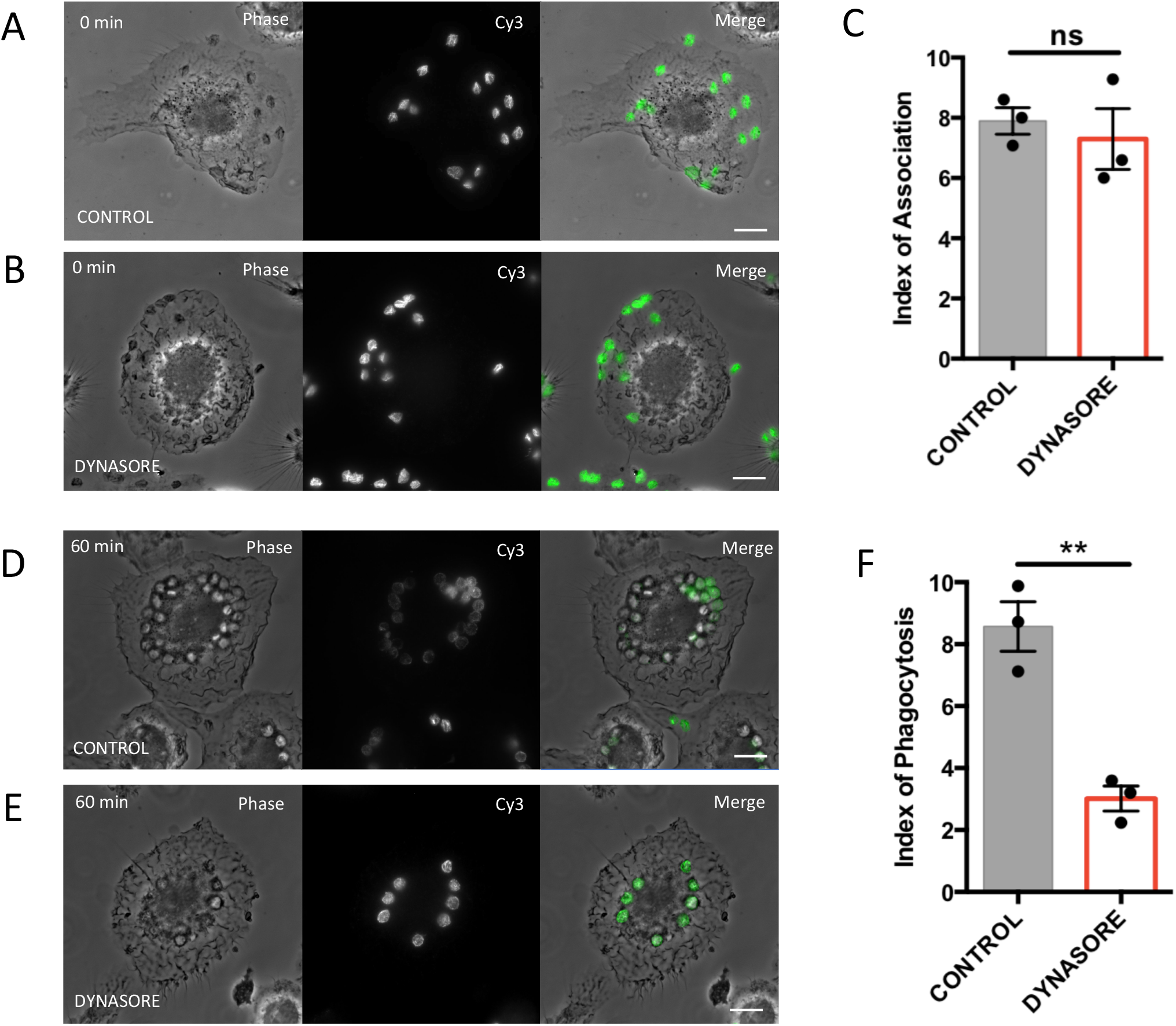
Pharmacological Inhibition of Dynamin-II impairs phagocytosis but not association. hMDMs were treated with DMSO (control, A & D) or Dynasore monohydrate (80 µm, B & E). hMDMs were then allowed to phagocytose IgM-C3 opsonised sRBC for 0 min (A & B) and 60 min (D & E), fixed and stained to detect sRBCs with Cy3-coupled anti-rabbit IgM antibodies. The indices of association (C) and phagocytosis (F) were calculated in 25 cells per donor at 0 and 60 min respectively. The means of 3 independent experiments are plotted (∗ p < 0,01). Bar = 10 μm.

### Inhibition of dynamin-2 decreases F-actin recruitment to the phagocytic cup

Phagocytosis is an active process requiring F-actin polymerisation to provide the force with which membranes are deformed to envelop the phagocytic target (Mularski and Niedergang, 2017; Torres-Gomez et al., 2020a). As CR3-mediated phagocytosis is impaired when dynamin-2 is inhibited, F-actin quantification was performed at the site of phagocytosis in cells treated with DMSO (control) or dynasore (80 μM). Phagocytosis was synchronised by centrifugation of hMDMs with C’-IgM-RBCs for 2 min. Macrophages were then allowed to phagocytose C’-IgM-RBCs for 15 min, then fixed and stained for external RBCs and F-actin. Quantification was performed on 4 different macrophage preparations from 4 different donors. In control cells, many F-actin rich phagocytic cups under C’-IgM-RBCs were easily located (Figure 3A). When dynamin-2 was inhibited, although sRBCs were associated with the membrane, there were less phagocytic cups (Figure 3B). Regions of interest were selected where C’-IgM-RBCs were attached and a phagocytic cup was visible. F-actin recruitment was quantified as described in the Materials and Methods section. The index of F-actin enrichment at sites of phagocytosis was 1.8±0.7 (Figure 3C) in the control cells treated with DMSO. Macrophages pre-treated with dynasore monohydrate exhibited a significantly lower F-actin ratio cup to body index: 0.75±0.4 (Figure 3D), representing a greater than 50% decrease in the F-actin recruitment at the site of CR3-phagocytosis in dynasore-treated macrophages. Therefore, F-actin enrichment in CR3 phagocytic cups is decreased when dynamin-2 is inhibited by dynasore monohydrate implicating dynamin-2 in the actin assembly of the CR3 phagocytic cup.

**Figure 3:**
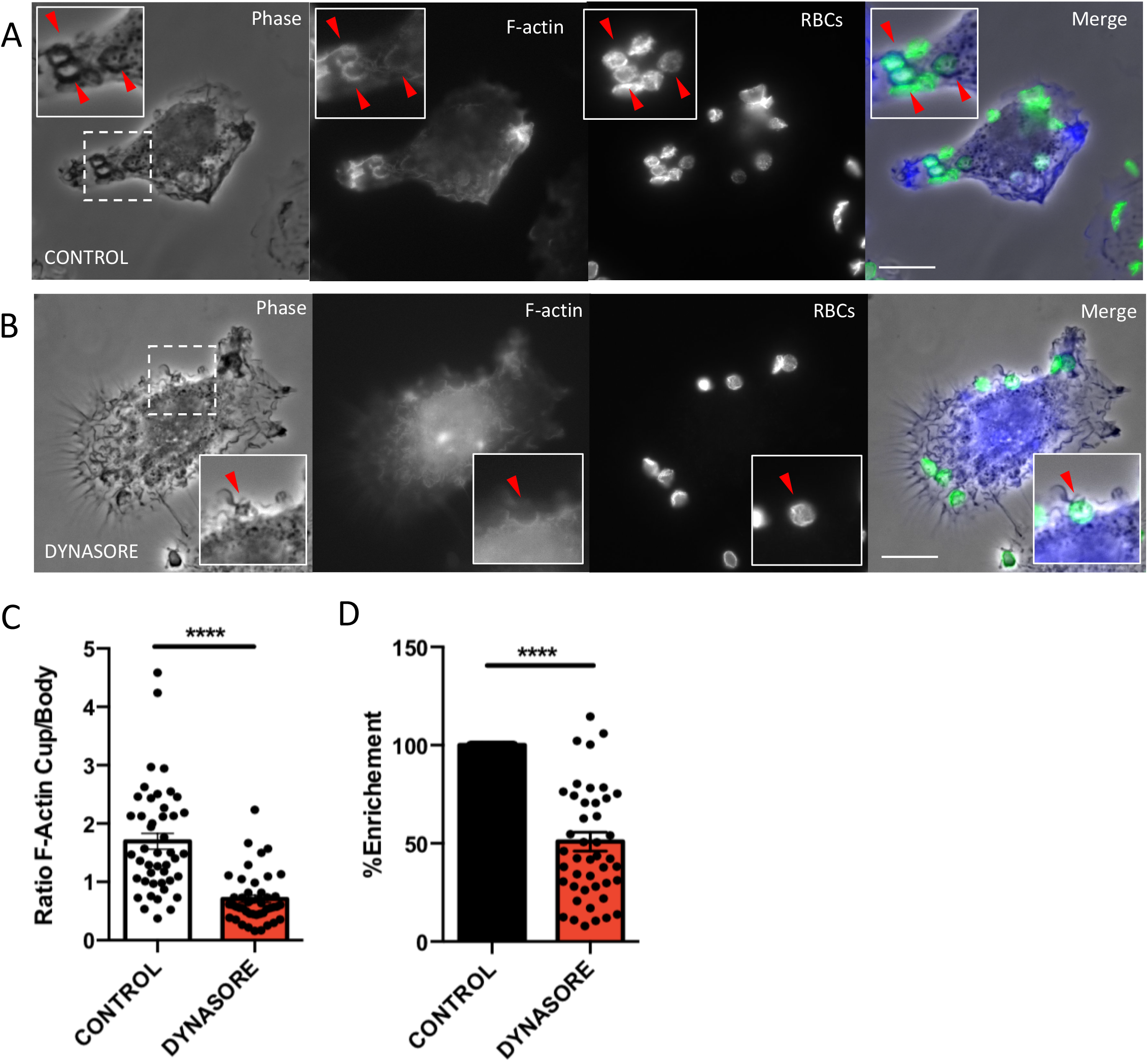
Pharmacological inhibition of dynamin-2 decreases the F-actin recruitment to the phagocytic cup. hMDMs were treated with DMSO (control) or dynasore monohydrate (80 µm). hMDMs were allowed to phagocytose IgM-C3 opsonised sRBC for 15 min, then fixed and stained to detect sRBCs with Cy3-coupled anti-rabbit IgM antibodies, as well as F-actin with phalloidin-Alexa488. Images for DMSO (A) and dynasore treated (B) cells were acquired using an inverted wide-field microscope (Leica DMI6000) with a 100× (1.4 NA) objective. Bar, 10 μm. Actin enrichment was quantified in phagocytic cups (A, B, red arrows), maximum fluorescence intensities of selected regions of interest were background corrected by subtracting the maximum intensity value from a membrane region in the same cell with no phagocytic event. Ratio values obtained by dividing the fluorescence intensities in the phagocytic cups by fluorescence intensities in the cell body were calculated (C). This ratio can be expressed as a percentage of F-actin enrichment relative to control cells (D). The results of four independent experiments are plotted (n = 15 actin cups/condition, ∗∗∗∗ p < 0.01). Bar = 10 μm, inset box = 12 μm.

### Timely inhibition of Dynamin-2 prevents CR3 phagosome scission in living cells

The role of Dynamin-2 in mediating scission of endocytic vesicles from the plasma membrane has been well characterised (Ferguson and De Camilli, 2012; Morlot and Roux, 2013). To determine if dynamin-2 is implicated in the last steps of phagosome closure in CR3-mediated phagocytosis, living RAW 264.7 macrophages transiently expressing Lifeact-mCherry and dynamin-2-GFP were allowed to start assembling a phagosome in the 3D phagosome formation and closure assay set-up (Marie-Anais et al., 2016a; Mularski et al., 2018). This mode of observation (shown schematically in Figure 4A) allows for simultaneous visualization of the phagocytic cup extension and closure with TIRF illumination, and the base of the phagocytic cup to be observed using epifluorescence (at an elevation of 3 μm above the coverslip). As macrophages neared the step of phagosome closure, the activity of dynamin was inhibited by the direct application of dynasore monohydrate to the phagocytosing cell. A representative cell is shown in Figure 4B. In the first 90 s of observation, prior to the addition of dynasore monohydrate, TIRF illumination reveals a cell undergoing phagosome closure, with concomitant accumulation of F-actin and dynamin-2 and a diminishing view of the RBC. After the addition of dynasore monohydrate at 90 s, a maximum in the F-actin and Dynamin-2 fluorescence is reached, followed by clearance of both proteins from the site without closure occurring (as the RBC is still visible). In the F-actin epifluorescence images, we see that prior to the addition of dynasore monohydrate, the RBC is in the ‘neck’ of the cell. The Dynamin-2 epifluorescence images indicate that Dynamin-2 is not well expressed in the base of the phagocytic cup, meaning that Dynamin-2 clears from the region before F-actin, though the position of the RBC is still visible. After the addition of dynasore monohydrate, the epifluorescence images indicate that the RBC has moved into the cell body, though the TIRFM images indicate that closure has not taken place. The epifluorescence images reveal that scission has not taken place. These experiments demonstrate that dynamin-2 accumulates together with F-actin and that timely inhibition of Dynamin-2 in living macrophages prevents CR3 phagosome closure and scission.

**Figure 4:**
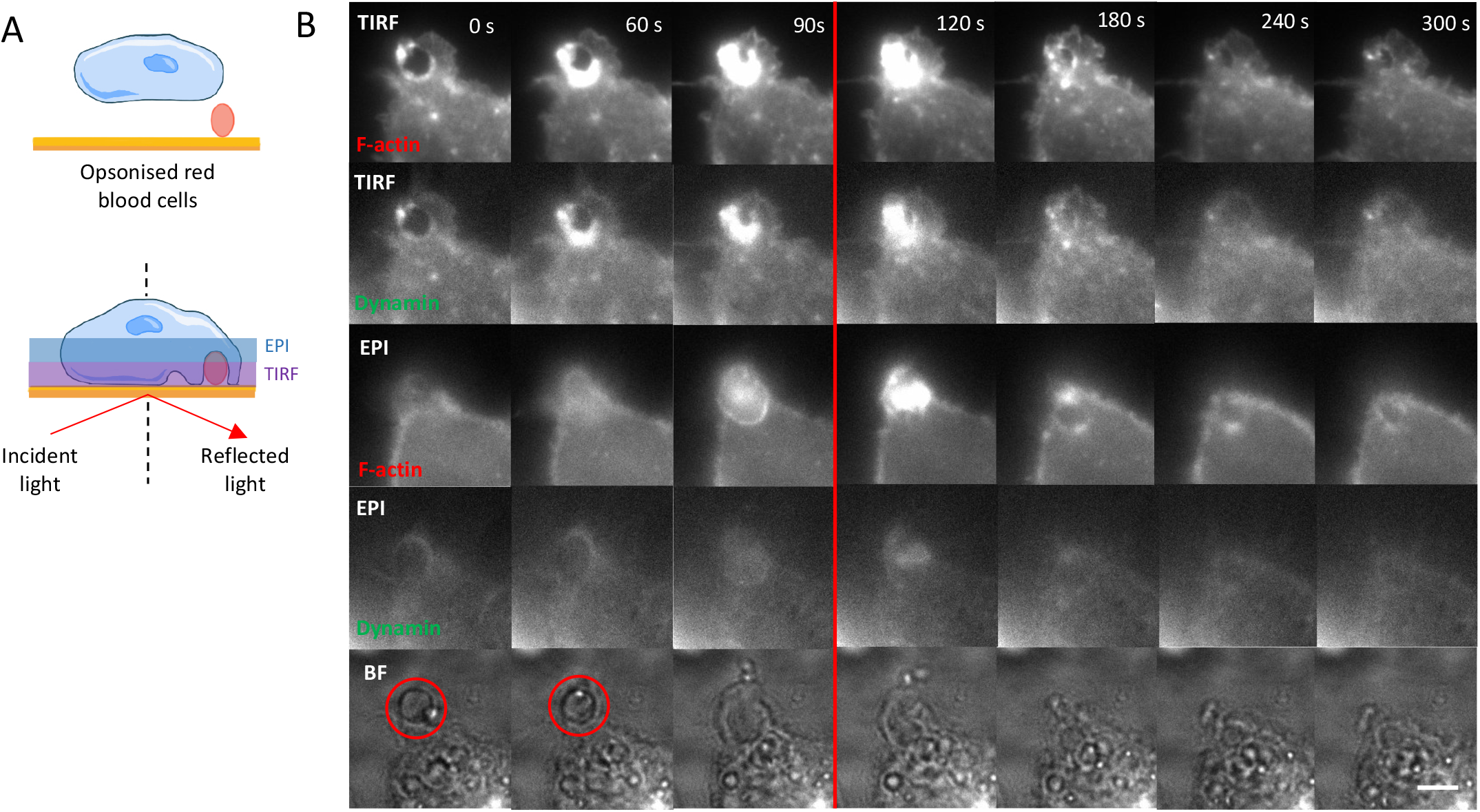
Inhibition of Dynamin-2 prevents phagosome scission in living cells. Phagosome closure assay was performed using RAW 264.7 macrophages transiently expressing both Lifeact-mcherry and Dynamin-2-GFP. Shown schematically in A, cells are allowed to phagocytose opsonized particles that are non-covalently bound to the coverslip, allowing the site of closure to be observed using TIRFM, and the base of the phagocytic cup to be observed using epifluorescence (at an elevation of 3 μm above the coverslip). TIRFM images of Lifeact-mcherry (B, top row) and Dynamin-2-GFP (B, second row), epifluorescence images of Lifeact-mcherry (B, third row) and Dynamin-2-GFP (B, fourth row) and brightfield images (B, bottom row) were captured every 2 s for 10 min at 37°. The RBC phagocytosed is clearly visible in brightfield images at 0 s and 60 s (B, bottom row, red circles), but pulled from the focal plane by the cell from 90s. At 90 s, Dynasore monohydrate was added to the fluid cell such that the final concentration was 80 μM (B, red line). The epifluorescence images show that the RBC has been internalised. Bar = 5 μm.

## Discussion

In this study, we reveal that dynamin-2 is crucial for CR3 integrin-mediated phagosome completion and closure. Dynamin-2 acts at two levels, both during actin polymerisation and membrane deformation, and during phagosome closure.

The CR3-mediated phagocytosis was often described as a “sinking” process relying on the formation of reduced phagocytic cups (Allen and Aderem, 1996; Kaplan et al., 1977). As pointed out recently (Jaumouille et al., 2019; Patel and Harrison, 2008), and when analysed using live cell imaging methods, it becomes visible that this type of phagocytosis does also generate some membrane extensions. dSTORM and live TIRFM experiments used in this study revealed ‘waves’ of F-actin that close over the phagocytic target (Figures 1 and 2), rather than an F-actin ring, characteristic of FcR-mediated phagocytosis, that closes around the target (Allen and Aderem, 1996; Marie-Anais et al., 2016b; Marion et al., 2012; Schlam et al., 2015). As CR3 is an integrin, formed by the αMβ2 pair, the actin structures formed around the target particle are similar to focal adhesions, in that they contain talin, vinculin and associated proteins (Torres-Gomez et al., 2020a). Integrins activity is highly regulated by conformational changes. During the “inside-out” phase of integrin activation, there is an actomyosin-dependent stretching of talin that allows the release of RIAM and subsequent binding to vinculin (Torres-Gomez et al., 2020b; Vigouroux et al., 2020), which reinforces the association to the β2 tail and its activation. Interestingly, dynamin-2 was not implicated in regulating these actin-dependent steps of integrin activation in human macrophages, as the association of macrophages to the complement-opsonised particles was not modified when dynamin-2 activity was perturbed (Figure 2). In the same line, the surface delivery of the CR3 receptors is not altered, as association of complement-opsonized RBCs was no different than in control cells when dynamin activity was impaired.

Instead, dynamin-2 was critical for the “outside-in” signal downstream of CR3 leading to actin polymerisation. This complex pathway involves the tyrosine kinase Syk, the small GTPases of the Rho family RhoG, and the actin nucleators VASP, Arp2/3 and Formin, as well as an actomyosin activity to orchestrate this integrin-mediated phagosome formation (Torres-Gomez et al., 2020a). Here we provide novel evidence that dynamin-2 controls actin polymerisation as well as membrane scission downstream of the phagocytic CR3. Indeed, dynamin-2 has been shown to interact directly with actin *in vitro*, and its interaction with short F-actin filaments was found to promote its GTPase activity and its oligomerisation, which was confirmed *in cellulo* (Gu et al., 2014). It was recently further deciphered how dynamin bundles actin filaments *in vitro* and how this mechanism could play a role in the context of cell fusion (Zhang et al., 2020). The assembly-stimulated GTPase activity of dynamin triggers rapid GTP hydrolysis and helix disassembly. This mechanism of actin bundling could also be relevant in the context of CR3-phagocytosis. We have indeed shown previously that the integrity of the F-actin network is necessary for sustained recruitment and function of dynamin-2 downstream of the phagocytic receptors for the Fc portion of Immunoglobulins (FcR) and, reversely, that dynamin-2 activity was necessary for an efficient assembly of the F-actin cup upon FcR ligation (Marie-Anais et al., 2016b). This is also in line with observations that actin polymerisation precedes dynamin recruitment at clathrin-coated pits in the last phases of clathrin-mediated endocytosis before the scission and release of vesicles (Grassart et al., 2014; Merrifield et al., 2002). Given the specificities of the CR3 integrin signaling and morphological features of the phagocytic cups, that dynamin-2 plays a crucial role during CR3-phagosome shaping and closing is particularly noteworthy.

Live cell imaging was instrumental to determine that dynamin-2 is required for CR3-phagosome closure and scission. Simultaneous TIRF and epifluorescence observation of living macrophages undergoing CR3 mediated phagocytosis allowed visualisation of dynasore monohydrate induced stalling of phagocytosis: the partially internalised phagocytic target still visible as dynamin-2 and F actin clear from the site. This result suggests that dynamin-2 is required for the membrane scission. Interestingly, the macropinocytic capture that shares many morphological features with CR3-phagocytosis has been described to be independent from dynamin activity (Le Roux et al., 2012; Liberali et al., 2008). In the context of CR3-phagocytosis, F-actin could help control the membrane tension and therefore complement the constriction activity of dynamin-2 for scission, as proposed (Morlot and Roux, 2013). Thus, dynamin might contribute to the mechano-sensing and –responding strategies used by the phagocytic cells to adapt their response to the physical properties of their targets, which is highly relevant to clear pathogens, debris and tumor cells (Mularski and Niedergang, 2017).

## Materials and Methods

### Antibodies, Plasmids and Reagents

The following antibodies were used: rabbit anti-Sheep RBCs (SRBC) (ICN Biochemicals); F(ab’)2-donkey anti-rabbit IgM (Jackson Immunoresearch); anti-dynamin-2 (kindly provided, via Mathieu Boissan (Hospital Saint-Antoine, Paris), by the De Camilli Lab (Ferguson et al., 2009)); F(ab’)2-Goat anti-Rabbit IgG-Alexa Fluor 555 (Life Technologies); C5 deficient serum human (Sigma); Phalloidin-Alexa 488 & 647 (Molecular Probes). The plasmid encoding Lifeact-mCherry was kindly provided by G. Montagnac (Institut Curie); the plasmids encoding DynWT-GFP was provided by Mark Mc Niven (Mayo Clinic). Dynasore monohydrate (Merck) was used at 80 µm in phagocytosis assays and in phagosome closure assays.

### Cell Culture

RAW264.7 macrophages were maintained in complete medium consisting of RPMI 1640-glutamax supplemented with 10 mM HEPES, 2 mM L-glutamine, 1 mM sodium pyruvate, 50 μM β-mercaptoethanol and 10% fetal calf serum (all from Gibco).

Human peripheral blood mononuclear cells were isolated from whole blood of healthy donors (Etablissement Français du Sang Ile-de-France, Site Trinité, INSERM agreement #18/EFS/030 ensures all donors have provided written informed consent to provide anonymous samples) by density gradient sedimentation using Ficoll-Plaque (GE Healthcare). Cells were allowed to adhere in 6 or 24 well plates at 37°C for 2 h in adhesion medium (RPMI 1640-Glutamax supplemented with 2 mM L-glutamine and 100 μg/ml streptomycin/penicillin) before cultivating them in complete medium (RPMI 1640-glutamax supplemented with 10%FCS, 2 mM L-glutamine and 100 μg/ml streptomycin/penicillin). Every 2 days, complete medium was exchanged. The adherent monocytes were left to differentiate into macrophages (referred to as human monocyte-derived macrophages (hMDM) in the text as described previously (Jubrail et al., 2020)and used for experiments from day 5 to day 10 of differentiation.

### Phagocytosis Assay

Phagocytosis Assays were performed with human monocyte-derived macrophages (hMDM) on 12 mm glass coverslips before incubation with complement bound, IgM-opsonized sheep red blood cells (C-IgM-RBC) as described elsewhere (Braun et al., 2004). C-IgM-RBC are resuspended in serum-free medium containing dynamin-2 inhibitors: dynasore monohydrate (80 μM) or DMSO as a control. Optimal dynasore monohydrate concentration was determined in previous studies (Marie-Anais et al., 2016b). hMDMs were exposed to inhibitors for 10 min prior to incubation with C-IgM-RBCs. After 2 min of centrifugation (500 g, RT) to ensure sedimentation of C-IgM-RBCs. Macrophages were allowed to internalise C-IgM-RBCs at 37 °C for specified times then fixed in 4% PFA (ParaFormAldehyde 4 %, PBS 1X) for 45 min at 4 °C. Cells were then permeabilised labelled with F(ab’)2 anti-rabbit IgG coupled to Cy3 in PBS1X/2% FCS before permeabilisation with 0.05% saponin and staining with phalloidin Alexa-488 in PBS1X/saponin 0.05%/2% FCS, and nuclei were stained with DAPI (DiAmidino Phenyl Indole).

External and internalised RBCs were counted in 25 randomly chosen cells, and the phagocytic index, i.e. the mean number of phagocytosed RBCs per cell, was calculated. The number of cell-associated (bound + internalised) RBCs, was also counted, to calculate the association index (mean number of associated RBCs per cell). Both phagocytic and attachment indices are expressed as percentages of control DMSO treated cells. Images were acquired using an inverted wide-field microscope (Leica DMI6000) with a 100× (1.4 NA) objective and a (MicroMAX Princeton Instruments) camera. Actin enrichment was quantified in phagocytic cups, maximum and average fluorescence intensities of a selected region of interest were background corrected by subtracting the mean value from a membrane region in the same cell with no phagocytosis event. Ratio values obtained by dividing the fluorescence intensities in the phagocytic cups by fluorescence intensities in the cell body were calculated as in (Braun et al., 2007).

### Electroporation of RAW264.7 cells

Cells were transfected by electroporation with Cell Project electrobuffer kits. One 100 mm plate of cells, grown to sub-confluence, and 20 μg of plasmid (10 μg per plasmid for co-transfection) were used for each electroporation. Cells were electroporated in 4 mm cuvettes (Biorad) at 250 V, 900 μF in an electroporation apparatus (X pulser Bio-Rad Laboratories) then immediately resuspended in complete culture medium and plated. Cells were used for experiments 15 – 18 hours post electroporation.

### Phagosome Closure Assay

Functionalised glass bottom dishes for phagosome closure assays were prepared according to the method described in (Marie-Anais et al., 2016a; Mularski et al., 2018). Briefly, C-IgM-SRBCs were centrifuged onto 35 mm glass bottom dishes (MatTek) pretreated with 0.01% poly-L-lysine in PBS for 30 min at RT. After washing once with 10% BSA in PBS, dishes were incubated in 10% BSA to saturate exposed poly-L-lysine. Dishes were then washed and filled with pre-warmed serum-free microscopy medium and maintained at 37 °C. Transfected cells were then allowed to sediment onto C-IgM-RBCs adhered to the dish surface.

### Total Internal Reflection Fluorescence Microscopy (TIRFM)

TIRFM was performed using a Till PHOTONICS iMIC microscope equipped with an oil‐immersion objective (Apo N 100×, NA1.49 Olympus America Inc.), a heating chamber maintained at 37°C and two cameras: a cooled iXonEM camera and an iXon3 897 Single Photon Detection EMCCD Camera (Andor Technology). The critical angle was determined prior to imaging by scanning through incident angles from 0° to 5° to maximise evanescent wave induced fluorescence (Marie-Anais et al., 2016a). Excitation was performed with 491 and 561 nm lasers. For the phagosome closure assay, images were acquired at 50 ms per frame in bright field illumination, TIRF mode and in epifluorescence mode with Polychrome illumination at 3 µm increment. TIRFM image stacks were processed using ImageJ Color Profiler software (NIH).

### Direct stochastic optical reconstruction microscopy (dSTORM)

Prior to fixaxion, coverslips were washed three times in PHEM buffer preheated to 37° C (PHEM buffer: 60mM PIPES, 25mM HEPES, 5mM EGTA, 2mM Mg (CH3CO) 2 (Sigma), pH6.9). Cells were then fixed for 15 min in 4% PFA and 0.02% glutaraldehyde (Sigma) preheated to 37° C. Permeabilization was then performed for 15 min at RT in 1X PBS / 4% PFA, 0.02% glutaraldehyde and 0.5% TritonX100 (Sigma). Cells were incubated with the polyclonal anti-dynamin-2, then with with F(ab’)2-Goat anti-Rabbit IgG-Alexa Fluor 555 (Life Technologies). Coverslips were mounted on microscope slides with 80μL cavities filled with optimized dSTORM buffer (Abbelight) containing phalloidin-647 (Molecular Probes) for F-actin staining. Microscopy was performed on a Leica SR GSD (DMI 6000B) equipped with an oil immersion objective (HCX PL APO 160x, NA1.43, Leica), 532 and 642 nm lasers and an EMCCD Andor iXon Ultra 897. Protein enrichment and localisation was quantified using ThunderSTORM (Ovesny et al., 2014) and SR-Tesseler (Levet et al., 2015). Tesselation based colocalization analysis was performed using Coloc-Tesseler software (Levet et al., 2015).

## Acknowledgements

The authors thank the IMAG’IC facility members and in particular Beatrice Durel for their help with dSTORM, and Christophe Le Clainche (Institute for Integrative Biology, Gif-sur-Yvette, France) for discussions on the project. IMAG’IC is a member of the National Infrastructure France BioImaging (ANR-10-INBS-04). The authors thank Sophie Echène for the illustrations in this paper.

